# DualLoc: Full-parameter fine-tuning of cascaded dual transformers for protein subcellular localization prediction

**DOI:** 10.64898/2026.03.27.714699

**Authors:** Yan Guang Chen, Wen-Yu Chung, Kuan Y. Chang

## Abstract

Accurate protein subcellular localization is essential for biological function, and mislocalization is linked to numerous diseases. While current methods like DeepLoc 2.0 employ lightweight fine-tuning of protein language models (PLMs), their ability to predict multi-compartment localization remains limited. To address this, we introduce DualLoc, a multi-label localization predictor for ten compartments. DualLoc leverages full-parameter fine-tuning of a cascaded dual-transformer architecture, built upon foundational PLMs and augmented with attention and dropout layers. We evaluated this framework using three foundational PLMs—ProtBERT, ESM-2, and ProtT5—as backbones. Cross-validation on Swiss-Prot and independent validation on the Human Protein Atlas demonstrate consistent superiority over state-of-the-art baselines. The best-performing variant, DualLoc-ProtT5, achieves 0.5872 accuracy, 0.8271 micro-F1, and 0.7811 macro-F1, with substantial gains in the Matthews correlation coefficient for the nucleus (+0.13), cell membrane (+0.13), and extracellular space (+0.07). Pointwise mutual information analysis of model outputs reveals biologically relevant compartment couplings, notably between the Golgi apparatus and endoplasmic reticulum (PMI = 0.25, P < 10^−6^), accurately reflecting secretory pathway coordination. DualLoc provides both a highly accurate predictive tool and a robust framework for investigating protein multi-localization mechanisms.

**Author summary:** Where a protein resides within a cell determines what it does. When proteins end up in the wrong location, normal cellular function breaks down—a misplacement linked to diseases like cancer and Alzheimer’s. While computational tools exist to predict these locations, accurately tracking proteins that multitask across multiple cellular compartments simultaneously remains a major challenge. We developed DualLoc, a new approach that predicts protein locations across ten different cellular compartments, from the nucleus to the cell membrane. By training an advanced artificial intelligence model on large protein sequence databases, our method more accurately identifies where proteins go, especially in complex, multi-location scenarios. Importantly, our analysis revealed meaningful biological patterns. We found strong predictive links between compartments that work closely together, such as the Golgi apparatus and the endoplasmic reticulum—two organelles that coordinate protein processing and transport. This suggests our model captures genuine cellular logic rather than simply memorizing data. By improving how we predict protein localization, DualLoc helps researchers better understand normal cellular function and disease mechanisms. Our method is freely available to the biomedical community.

## Introduction

In eukaryotic cells, proteins execute specific biological functions within distinct subcellular compartments, and their activity critically depends on accurate localization [1, 2]. Mislocalization can disrupt function, leading to pathological conditions such as metabolic disorders, cardiovascular and neurodegenerative diseases, and cancer [2, 3]. Consequently, accurately predicting protein subcellular localization has emerged as a vital focus in bioinformatics research [4–7].

Historically, subcellular localization prediction relied primarily on sequence features analyzed using machine learning. Early tools such as YLoc [8] pioneered interpretable probabilistic models that integrate features such as sorting signals and transmembrane segments and provide prediction confidence scores. Subsequent advancements included DeepLoc [6], which introduced an end-to-end learning approach using bidirectional long short-term memory networks for raw amino acid sequences, and BUSCA [9], which improved eukaryotic prediction accuracy by combining multiple specialized tools via a weighted decision strategy. Ongoing evolution has yielded tools such as SCLpred-ECL [10], which employs deep N-to-1 convolutional neural networks with ensemble learning to simultaneously predict across eight major compartments, and LocPro [11], which integrates convolutional and long short-term memory networks to improve sequence-based prediction.

Recent major progress has been driven by applying transformer architectures [12] to protein sequence analysis. These models leverage inherent attention mechanisms to capture both local amino acid patterns and global structural features, enabling the development of large-scale protein language models (PLMs) like ESM [13] and ProtTrans [14]. Such PLMs undergo unsupervised pretraining on vast unlabelled sequence datasets to generate high-dimensional embeddings rich in biophysical meaning. The resulting frozen models can then be efficiently fine-tuned for downstream tasks like subcellular localization prediction, significantly boosting performance.

Tools exemplifying this approach include DeepLoc 2.0 [7], which combines ESM-1b [13] and ProtT5 [14] embeddings with attention layers and multilayer perceptrons; DeepLoc 2.1 [15], which enhances membrane protein classification accuracy by 8.2% to 12.7% using attention; and ProStructNet [16], which synergizes ProtT5 sequence embeddings with AlphaFold2-predicted [17] structural embeddings via a graph attention network and multilayer perceptron, achieving a further 4% gain in multilabel localization accuracy; and recent methodologies employing ESM-2 [18] feature refinement through Res-VAE/UMAP dimensionality reduction or Low Rank Adaptation (LoRA) fine-tuning of ESM-2/ProtT5. While these fine-tuned PLM-based approaches enhance performance with reduced computational demands, the extent to which they capture the full complexity of subcellular biology remains unclear and warrants investigation [19–22].

This study systematically evaluates the efficacy of full-parameter fine-tuning strategies for subcellular localization and signal type prediction using large-scale PLMs.

We selected three pretrained base models—ProtT5 [14], ESM-2 [18] (both high-accuracy), and the less explored ProtBERT [14]—each implemented as a cascaded dual-encoder architecture. Within the DualLoc framework, the architecture uses two distinct encoders, trained simultaneously, to capture multi-scale protein features. The first encoder is initialized with pretrained weights to leverage broad evolutionary and biological context, while the second is randomly initialized to specialize in learning task-specific patterns from scratch. By jointly optimizing both encoders, DualLoc effectively integrates global biological knowledge with high-resolution localization features. This cascaded dual-encoder system is further enhanced with a multi-head attention mechanism and dropout layers to optimize classification across ten subcellular compartments.

## Materials and methods

### Subcellular Localization Datasets

This study used the eukaryotic protein subcellular localization dataset from DeepLoc [7], derived from manually reviewed Swiss-Prot entries in UniProt release 2021 03 [23]. As curated by DeepLoc 2.0, the dataset exclusively contains experimentally validated (ECO:0000269) nuclear-encoded eukaryotic proteins with≥ 40 amino acids, excluding bacterial/archaeal sequences and protein fragments. The final curated dataset comprised 28,303 unique sequences categorized into 10 subcellular compartments: cytoplasm, nucleus, extracellular, cell membrane, mitochondrion, plastid, endoplasmic reticulum (ER), lysosome/vacuole, Golgi apparatus, and peroxisome S1 Table.

For independent validation, we employed Human Protein Atlas (HPA) data curated by DeepLoc 2.0 [7]. To ensure label reliability, only annotations with “Enhanced” or “Supported” confidence levels were retained. Proteins sharing *>*30% global sequence identity with the Swiss-Prot training set were excluded to mitigate homology bias, yielding 1,717 non-redundant sequences classified into eight compartments (excluding extracellular and plastid) consistent with the training set categories S2 Table.

### Sorting Signal Dataset

A high-confidence sorting signal dataset was compiled from annotations in DeepLoc 2.0 and Swiss-Prot [7, 23]. This dataset encompasses nine distinct sorting signal classes: signal peptides (SP), transmembrane segments (TM), mitochondrial targeting (MT) signals, chloroplast transit peptides (CH), thylakoid transfer peptides (TH), nuclear localization signals (NLS), nuclear export signals (NES), peroxisomal targeting signals (PTS), and glycosylphosphatidylinositol (GPI)-anchor signals S3 Table.

### Large Language Models (LLMs)

Three Transformer-based large language models—ProtBERT [14], ESM-2 [18], ProtT5 [14]—were employed for protein subcellular localization prediction. These models exhibit distinct architectures, pre-training data sources, and masking strategies that influence their capabilities [24, 25]. ProtBERT extends this with a 16-layer encoder trained on the Big Fantastic Database (BFD), demonstrating protein-specific semantic improvements at scale. ESM-2 advances sequence modeling with a 33-layer encoder that incorporates relative position bias to enhance long-sequence processing and structural inference. ProtT5 utilizes a 24-layer encoder-decoder framework with span masking to preserve global context while capturing local features and long-range interactions.

Model performance was evaluated using 5-fold cross-validation [26]. All models were fine-tuned for 20 epochs with early stopping when loss convergence was reached, followed by a final assessment on the independent test set.

### Design and Architecture of DualLoc

The DualLoc framework comprises two interacting transformer paths, jointly optimized via full-parameter fine-tuning (Fig 1). During inference, the first predicts subcellular localization and passes its learned representations to the second, which predicts sorting signals. Each path is built from cascaded dual PLMs—one pretrained and one randomly initialized.

**Fig 1.**
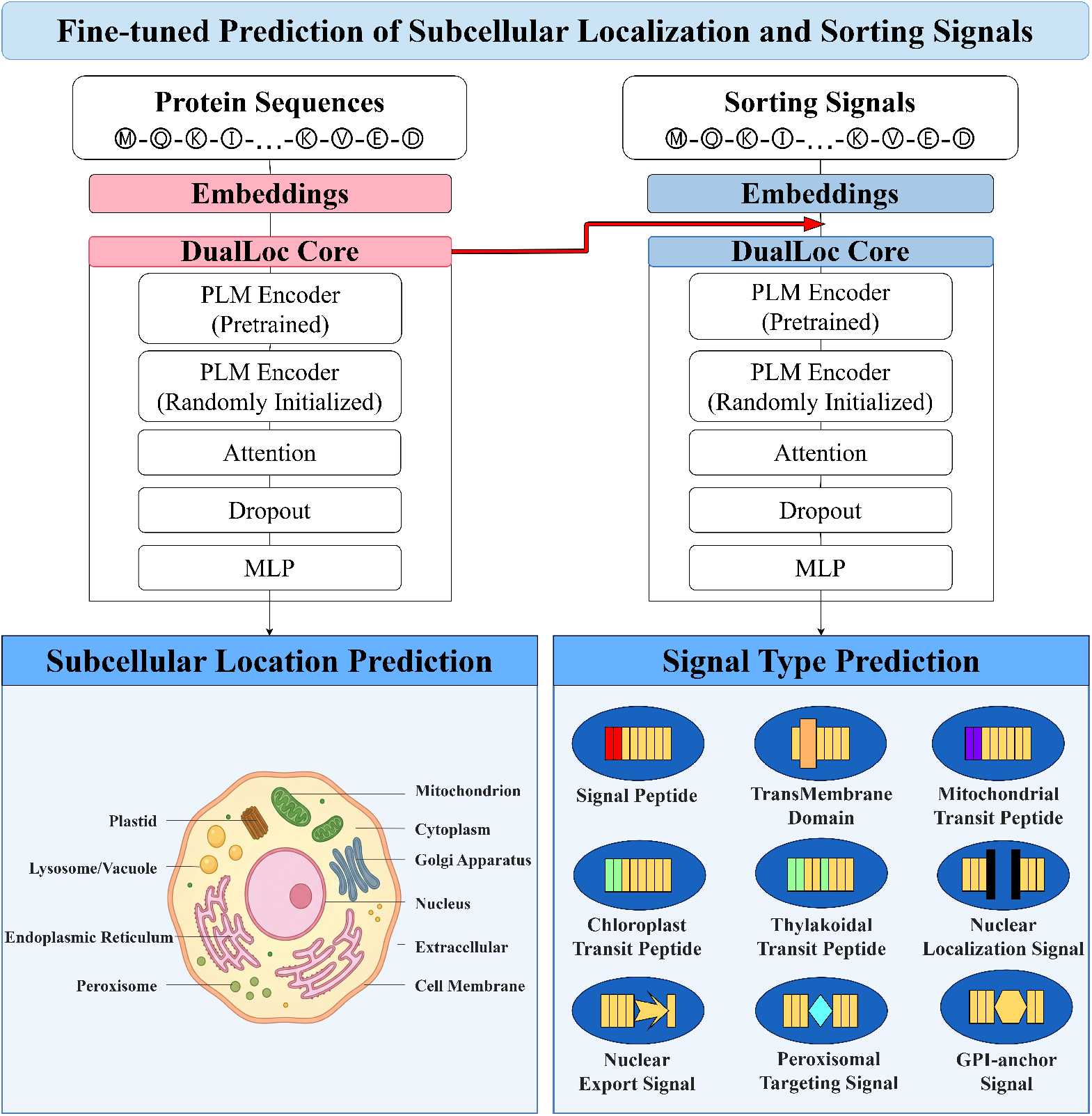
DualLoc Architecture Overview. Schematic representation of the DualLoc framework for sequential prediction of subcellular localization and protein sorting signals. An input amino acid sequence is encoded using one of three Transformer-based PLMs (ProtBERT, ESM-2, or ProtT5). The resulting embeddings are first used to predict ten subcellular localization labels. Features from this localization module are then fed into a parallel architecture that predicts nine sorting signal classes, as indicated by the red arrow. Both hierarchical stages utilize the cascaded dual PLM encoders, attention mechanisms, and dropout layers.

In the primary pathway, DualLoc performs end-to-end fine-tuning of both pre-trained and randomly initialized PLM embeddings within a cascaded architecture, enabling the model to capture intricate, non-linear dependencies within protein sequences. For an input protein sequence s= (x_1_,…,x_*n*_), the pre-trained encoder first generates an embedding matrix *E* ∈*R*^*n×d*^, where *n* is the sequence length and *d* is the hidden dimension, followed by average pooling and extraction of the 1024-dimensional [CLS] token feature vector (h_*CLS*_ ∈*R*^1024^). This vector is processed through an attention module (computing smoothed scores) [7], a dropout layer, and a linear layer, reducing dimensionality to 10 to predict localization probabilities. The pathway is optimized via binary cross-entropy loss, with full-parameter updates applied to both the cascaded dual PLMs and the attention, dropout, and linear layers during fine-tuning.

In the auxiliary pathway, DualLoc processes hidden representations from the primary pathway and integrates signal-specific embeddings. The [CLS] vector (h_*CLS*_ ∈ *R*^1024^) is concatenated with the 10-dimensional localization probabilities to form a 1034-dimensional input. This is processed through an isomorphic module (attention, dropout, linear layer), reducing dimensionality to 9 for sorting signal prediction, trained similarly with constrained backpropagation.

An integration layer subsequently fuses localization features with sorting signal probabilities to generate joint annotations for both protein properties. For both pathways, 20% of the training data served as a validation set for hyperparameter optimization and performance monitoring.

### Pointwise Mutual Information

Pointwise Mutual Information (PMI) [27] measures the strength of co-occurrence of a specific label pair (x, y) at the sample level. Its calculation formula is:

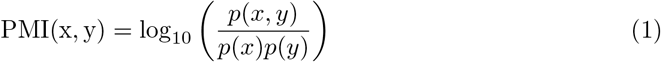

where p(x) denotes the probability of event x occurring, and p(x, y) is the joint probability of x and y occurring together. A positive value indicates that the pair co-occurs more frequently than expected under independence; a negative value indicates that it co-occurs less frequently than expected. This allows PMI to intuitively reflect the relative increase or decrease in co-occurrence strength compared to the independent case.

### Data visualization of subcellular localization protein sequences

To visualize subcellular localization patterns, protein sequence embeddings were extracted using four pre-trained language models, both before and after fine-tuning. Each 1024-dimensional representation vector was reduced to two dimensions via principal component analysis (PCA) [28], uniform manifold approximation and projection (UMAP) [29], and t-distributed stochastic neighbor embedding (t-SNE) [30] for spatial visualization. This dimensionality reduction revealed label-specific distribution patterns and quantified the discriminative capacity of embeddings for localization features, providing insights to optimize downstream classification models.

## Results

### Core Characteristics of Subcellular Localization Distribution

Analysis of protein localization patterns across ten subcellular compartments revealed distinct distribution characteristics (Fig 2). Proteins with single localizations are predominantly localized to the cytoplasm, nucleus, cell membrane, extracellular space, and mitochondria, collectively accounting for 82.4% of singly localized proteins. Among dual-localized proteins, the nucleus + cytoplasm combination was most frequent (60.3%), followed by nucleus + cell membrane (8.1%) and cell membrane + lysosome/vacuole (3.2%). Multilocalized proteins (≥3 compartments) were frequently found in the nucleus (59.1%), with the nucleus + cytoplasm + cell membrane triplet being the most common (17.9% of triple-localized proteins). Rare compartments, such as plastids and peroxisomes, primarily exhibited single- or dual-localization patterns (*>* 90%).

**Fig 2.**
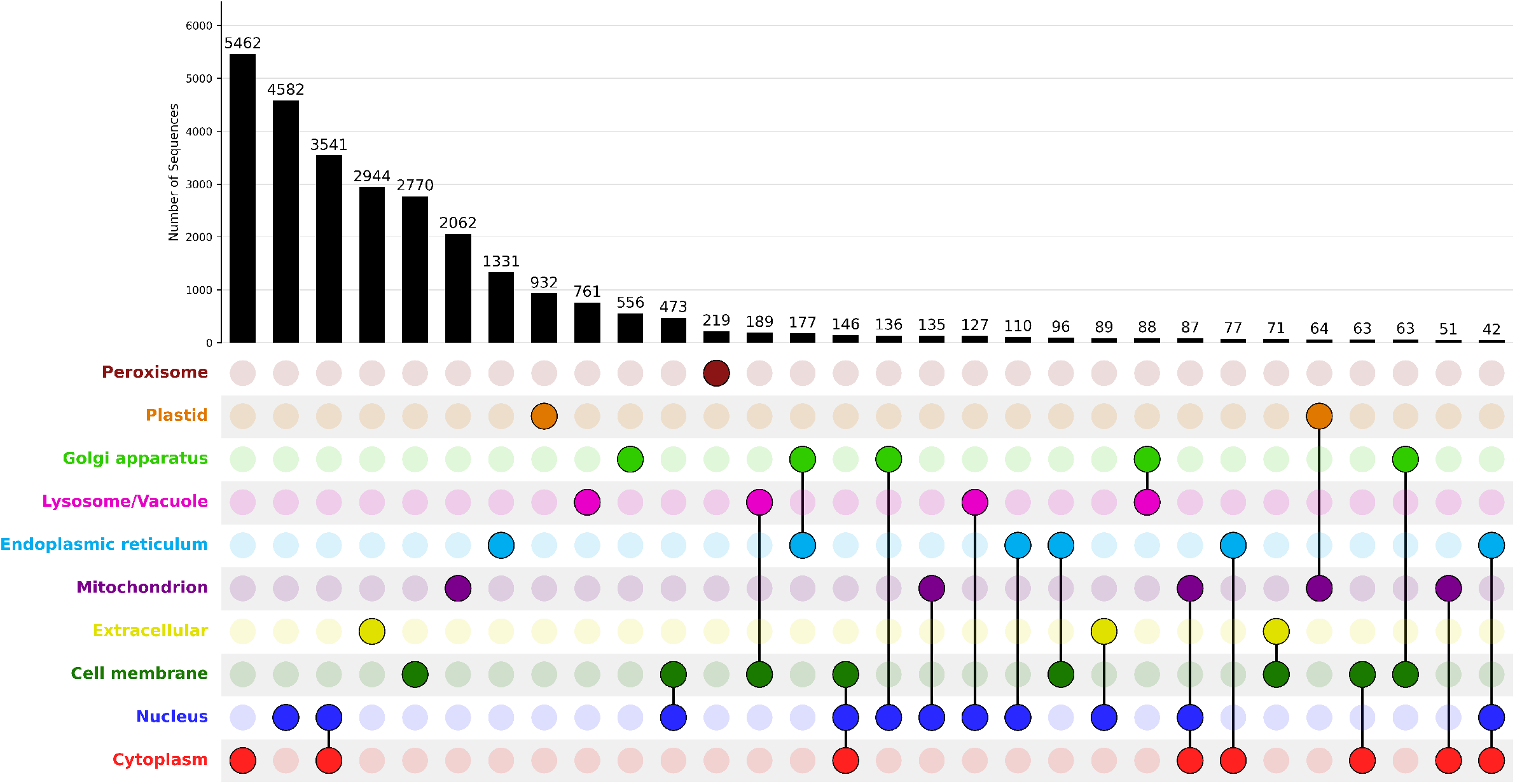
Most frequent subcellular localizations in the Swiss-Prot Dataset.

PMI analysis identified significant cooperative associations between compartments (Fig 3). The Golgi apparatus showed strong co-occurrence with both the ER (PMI = 0.25, p *<* 10^*−*6^) and the lysosome/vacuole (PMI = 0.11, p *<* 10^*−*6^).

**Fig 3.**
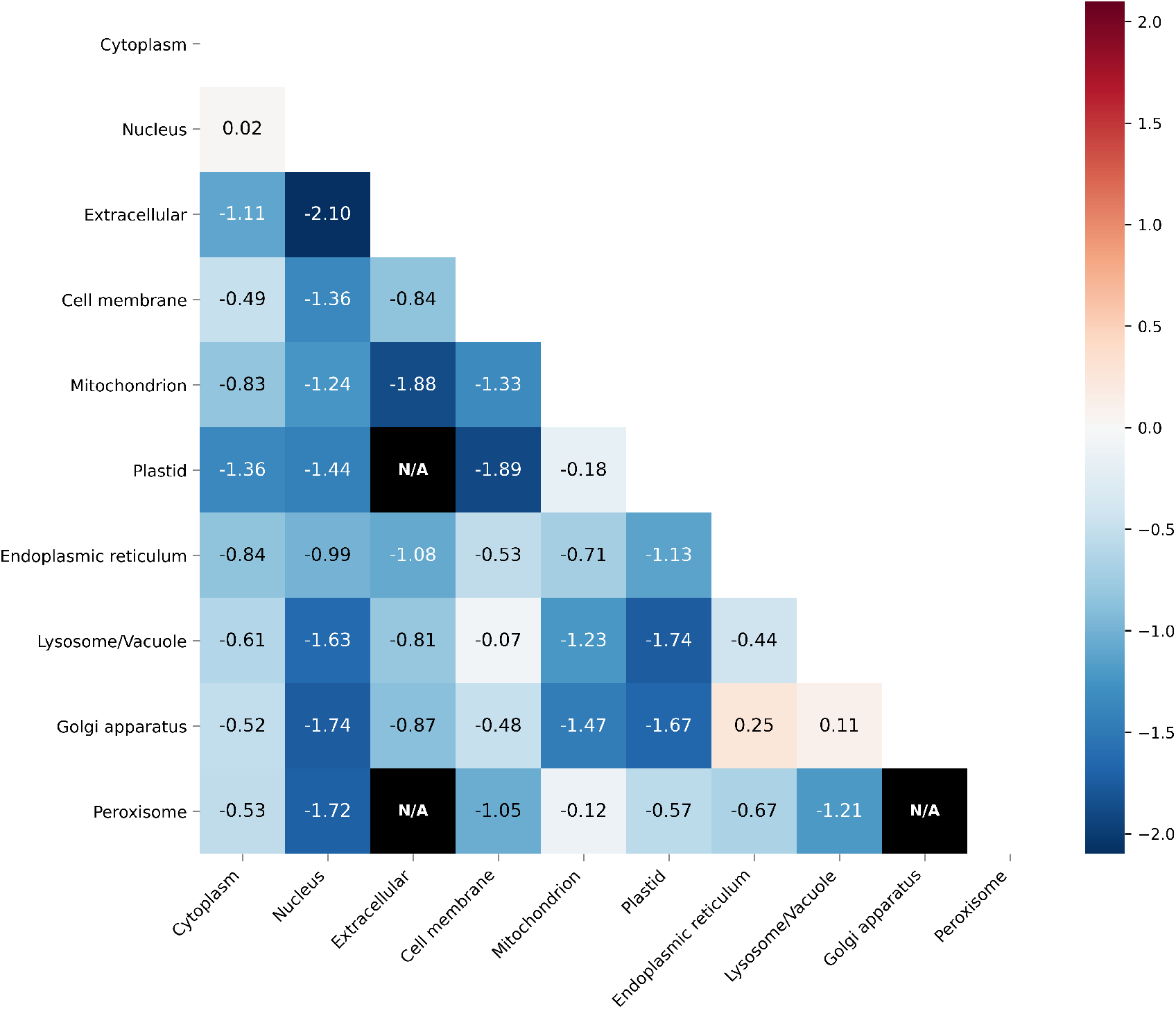
Co-occurrence analysis of subcellular compartments using Swiss-Prot localization data.

A Sankey diagram mapped nine sorting signals to subcellular compartments, illustrating how targeting signals direct proteins to single or multiple destinations (Fig 4). SP was present in 41.1% of proteins, with 80.2% of SP-bearing proteins localizing to the extracellular space, consistent with its role in the secretory pathway.

**Fig 4.**
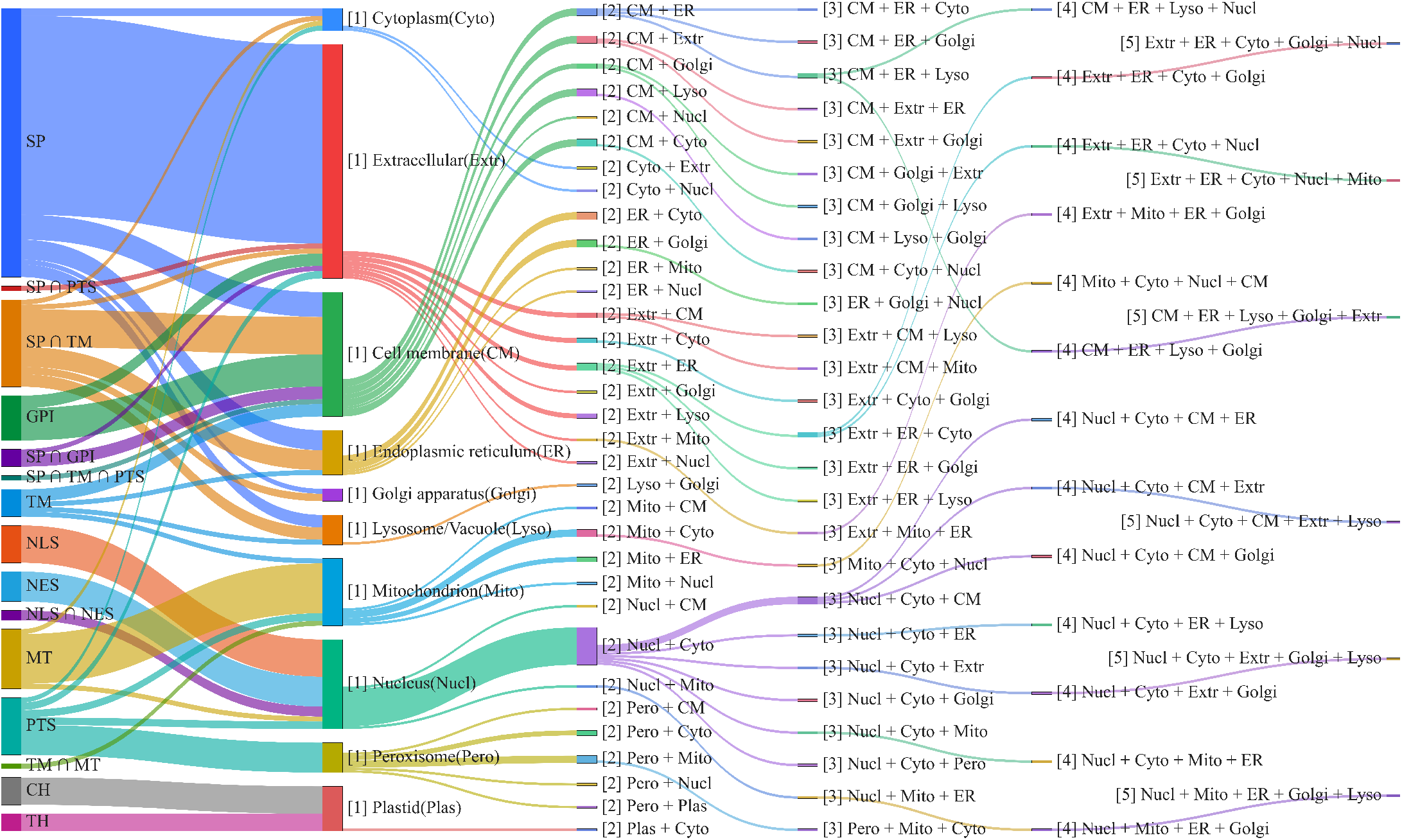
Protein Sorting Signal–Associated Subcellular Localization Flow.

TM primarily directed proteins to the cell membrane (69.9%). NLS and MT signals exhibited near-perfect specificity, localizing 100.0% and 99.2% of proteins to the nucleus and mitochondria, respectively. Analysis of dual-localization patterns revealed that combinations originating from the cell membrane or ER were most prominent. Proteins with NLS/NES were exclusively nuclear, while nucleus + cytoplasm combinations showed the greatest potential for triple localization.

Dimensionality reduction (PCA, UMAP, and t-SNE) of features from the non-fine-tuned ProtT5 model showed substantial overlap across localization classes S1A Fig. After fine-tuning ProtT5 on localization data, the separation in feature space improved significantly S1B Fig. While high-frequency compartments, such as the cytoplasm and nucleus, exhibited strong global separation, rare compartments formed distinct t-SNE clusters, demonstrating the model’s proficiency in resolving fine-grained localization features.

### Multilocalization Classification

We developed DualLoc, a novel model that integrates a dual-encoder architecture comprising one pretrained PLM and one randomly initialized large-scale PLM (ProtBERT, ESM-2, or ProtT5). Inspired by DeepLoc 2.0, DualLoc incorporates attention layers from the pretrained model, one-dimensional Gaussian smoothing to optimize attention weights, dropout layers, and fully connected classification layers. This architecture enhances the capture of both local signal motifs and long-range amino acid dependencies through comprehensive fine-tuning. In Swiss-Prot cross-validation for subcellular localization, DualLoc outperformed both a general-purpose BERT model and DeepLoc 2.0 (Table 1). Among four DualLoc variants, DualLoc-ProtT5 achieved the highest overall performance (Accuracy: 0.5872; Jaccard: 0.7812; micro-F1: 0.8371; macro-F1: 0.7811) S4 Table S5 Table S6 Table, confirming its effectiveness across both high and low-frequency localization labels. All variants showed improvements, validating the strategy for multilabel classification. Fine-grained analysis revealed compartment-specific strengths: DualLoc-ProtBERT excelled in cytoplasm prediction (MCC: 0.8561), DualLoc-ESM-2 led in lysosome/vacuole (MCC: 0.4631), and DualLoc-ProtT5 achieved the highest performance in nucleus (MCC: 0.8209), extracellular (MCC: 0.9221), and cell membrane (MCC: 0.7951), averaging a 2% MCC gain over the next best model.

**Table 1.** Performance Evaluation on the Swiss-Prot Dataset.

|  | DualLoc |  |  |  |
| --- | --- | --- | --- | --- |
|  | ProtBERT | ESM1b | ESM-2 | ProtT5 |
| Pred. Num. Labels (1.27) | 1.2387 ± 0.0152 | 1.2593 ± 0.0301 | 1.2675 ± 0.0224 | 1.2701 ± 0.0139 |
| Accuracy | 0.4528 ± 0.0512 | 0.5318 ± 0.0361 | 0.5347 ± 0.0314 | **0.5872 ± 0.0223** |
| Jaccard | 0.7382 ± 0.0081 | 0.7627 ± 0.0092 | 0.7713 ± 0.0118 | **0.7812 ± 0.0074** |
| MicroF1 | 0.8054 ± 0.0053 | 0.7854 ± 0.0029 | 0.8063 ± 0.0054 | **0.8371 ± 0.0056** |
| MacroF1 | 0.6762 ± 0.0075 | 0.6812 ± 0.0031 | 0.7502 ± 0.0067 | **0.7811 ± 0.0077** |
| MCC per location |  |  |  |  |
| – Cytoplasm | **0.8561 ± 0.0150** | 0.6440 ± 0.0073 | 0.7542 ± 0.0101 | 0.7695 ± 0.0111 |
| – Nucleus | 0.6975 ± 0.0098 | 0.7399 ± 0.0097 | 0.7756 ± 0.0072 | **0.8209 ± 0.0065** |
| – Extracellular | 0.7556 ± 0.0089 | 0.9111 ± 0.0092 | 0.9138 ± 0.0069 | **0.9221 ± 0.0070** |
| – Cell membrane | 0.7191 ± 0.0146 | 0.7180 ± 0.0070 | 0.7382 ± 0.0092 | **0.7951 ± 0.0096** |
| – Mitochondrion | 0.8373 ± 0.0207 | 0.8326 ± 0.0094 | 0.8452 ± 0.0107 | **0.8556 ± 0.0129** |
| – Plastid | 0.8992 ± 0.0212 | 0.9172 ± 0.0181 | 0.9174 ± 0.0143 | **0.9327 ± 0.0103** |
| – Endoplasmic reticulum | 0.6517 ± 0.0249 | 0.6976 ± 0.0172 | 0.6977 ± 0.0169 | **0.7133 ± 0.0161** |
| – Lysosome / Vacuole | 0.3372 ± 0.0511 | 0.3163 ± 0.0313 | **0.4631 ± 0.0137** | 0.3973 ± 0.0212 |
| – Golgi apparatus | 0.3970 ± 0.0283 | 0.4013 ± 0.0109 | 0.4879 ± 0.0196 | **0.5363 ± 0.0191** |
| – Peroxisome | 0.5473 ± 0.0347 | 0.4716 ± 0.0614 | 0.5757 ± 0.0415 | **0.6416 ± 0.0396** |
|  | DeepLoc 2.0 |  | ProStructNet |  |
|  | ESM1b | ProtT5 | ProtT5 |  |
| Pred. Num. Labels (1.27) | 1.27 ± 0.02 | 1.26 ± 0.02 | 1.27 ± 0.02 |  |
| Accuracy | 0.53 ± 0.02 | 0.55 ± 0.02 | **0.59 ± 0.02** |  |
| Jaccard | 0.68 ± 0.01 | 0.69 ± 0.01 | 0.73 ± 0.02 |  |
| MicroF1 | 0.72 ± 0.01 | 0.73 ± 0.01 | 0.75 ± 0.01 |  |
| MacroF1 | 0.64 ± 0.01 | 0.66 ± 0.01 | 0.69 ± 0.02 |  |
| MCC per location |  |  |  |  |
| – Cytoplasm | 0.61 ± 0.01 | 0.62 ± 0.01 | 0.65 ± 0.01 |  |
| – Nucleus | 0.66 ± 0.02 | 0.69 ± 0.01 | 0.70 ± 0.01 |  |
| – Extracellular | 0.85 ± 0.03 | 0.85 ± 0.04 | 0.88 ± 0.04 |  |
| – Cell membrane | 0.64 ± 0.01 | 0.66 ± 0.01 | 0.69 ± 0.01 |  |
| – Mitochondrion | 0.73 ± 0.03 | 0.76 ± 0.02 | 0.79 ± 0.02 |  |
| – Plastid | 0.88 ± 0.01 | 0.90 ± 0.01 | 0.92 ± 0.01 |  |
| – Endoplasmic reticulum | 0.52 ± 0.01 | 0.56 ± 0.03 | 0.62 ± 0.04 |  |
| – Lysosome / Vacuole | 0.24 ± 0.03 | 0.28 ± 0.04 | 0.31 ± 0.04 |  |
| – Golgi apparatus | 0.36 ± 0.06 | 0.34 ± 0.05 | 0.44 ± 0.06 |  |
| – Peroxisome | 0.48 ± 0.05 | 0.56 ± 0.08 | 0.58 ± 0.07 |  |

The best results are highlighted in bold.

Independent validation on the HPA confirmed DualLoc-ProtT5’s superior localization prediction, outperforming DeepLoc 2.0 across all key metrics (Accuracy: 0.4098; Jaccard: 0.5457; micro-F1: 0.6175; macro-F1: 0.5024; Table 2).

**Table 2.**
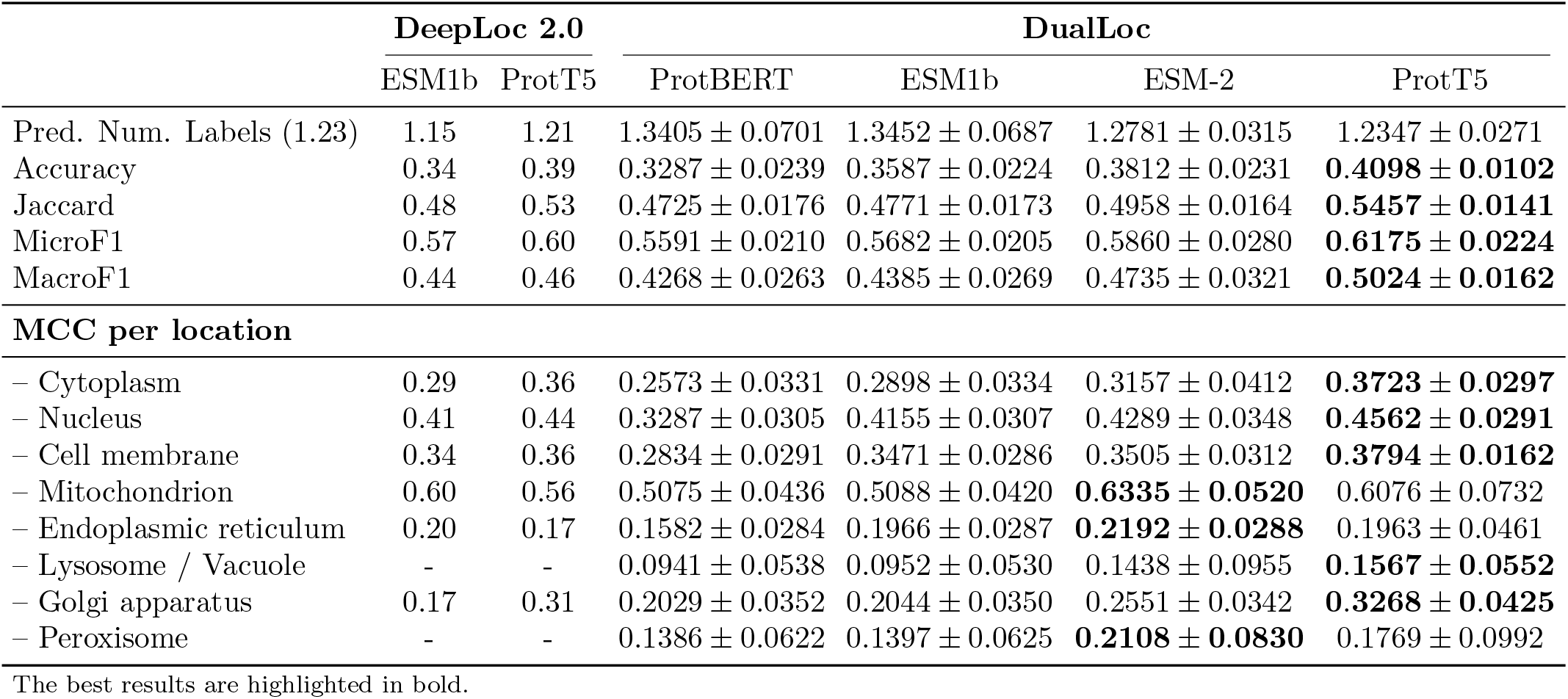
Performance Evaluation on the independent HPA Dataset.

|  | DeepLoc 2.0 |  | DualLoc |  |  |  |
| --- | --- | --- | --- | --- | --- | --- |
|  | ESM1b | ProtT5 | ProtBERT | ESM1b | ESM-2 | ProtT5 |
| Pred. Num. Labels (1.23) | 1.15 | 1.21 | $1.3405 \pm 0.0701$ | $1.3452 \pm 0.0687$ | $1.2781 \pm 0.0315$ | $1.2347 \pm 0.0271$ |
| Accuracy | 0.34 | 0.39 | $0.3287 \pm 0.0239$ | $0.3587 \pm 0.0224$ | $0.3812 \pm 0.0231$ | **$0.4098 \pm 0.0102$** |
| Jaccard | 0.48 | 0.53 | $0.4725 \pm 0.0176$ | $0.4771 \pm 0.0173$ | $0.4958 \pm 0.0164$ | **$0.5457 \pm 0.0141$** |
| MicroF1 | 0.57 | 0.60 | $0.5591 \pm 0.0210$ | $0.5682 \pm 0.0205$ | $0.5860 \pm 0.0280$ | **$0.6175 \pm 0.0224$** |
| MacroF1 | 0.44 | 0.46 | $0.4268 \pm 0.0263$ | $0.4385 \pm 0.0269$ | $0.4735 \pm 0.0321$ | **$0.5024 \pm 0.0162$** |
| **MCC per location** |  |  |  |  |  |  |
| – Cytoplasm | 0.29 | 0.36 | $0.2573 \pm 0.0331$ | $0.2898 \pm 0.0334$ | $0.3157 \pm 0.0412$ | **$0.3723 \pm 0.0297$** |
| – Nucleus | 0.41 | 0.44 | $0.3287 \pm 0.0305$ | $0.4155 \pm 0.0307$ | $0.4289 \pm 0.0348$ | **$0.4562 \pm 0.0291$** |
| – Cell membrane | 0.34 | 0.36 | $0.2834 \pm 0.0291$ | $0.3471 \pm 0.0286$ | $0.3505 \pm 0.0312$ | **$0.3794 \pm 0.0162$** |
| – Mitochondrion | 0.60 | 0.56 | $0.5075 \pm 0.0436$ | $0.5088 \pm 0.0420$ | **$0.6335 \pm 0.0520$** | $0.6076 \pm 0.0732$ |
| – Endoplasmic reticulum | 0.20 | 0.17 | $0.1582 \pm 0.0284$ | $0.1966 \pm 0.0287$ | **$0.2192 \pm 0.0288$** | $0.1963 \pm 0.0461$ |
| – Lysosome / Vacuole | - | - | $0.0941 \pm 0.0538$ | $0.0952 \pm 0.0530$ | $0.1438 \pm 0.0955$ | **$0.1567 \pm 0.0552$** |
| – Golgi apparatus | 0.17 | 0.31 | $0.2029 \pm 0.0352$ | $0.2044 \pm 0.0350$ | $0.2551 \pm 0.0342$ | **$0.3268 \pm 0.0425$** |
| – Peroxisome | - | - | $0.1386 \pm 0.0622$ | $0.1397 \pm 0.0625$ | **$0.2108 \pm 0.0830$** | $0.1769 \pm 0.0992$ |

The best results are highlighted in bold.

Compartment-specific analysis showed that DualLoc-ProtT5 achieved the highest performance across five compartments, including the cytoplasm (MCC: 0.3190) and the nucleus (MCC: 0.4562). Conversely, DualLoc-ESM-2-based variants led in mitochondria (MCC: 0.6335), ER (MCC: 0.2192), and peroxisome (MCC: 0.2108). This performance variation across models and compartments demonstrates that the dual-encoder pretraining architecture effectively captures compartment-specific biological signatures, enhancing multilabel classification capability and improving generalizability across datasets

### Classification of Protein Sorting Signals

In sorting signals classification, DualLoc achieved significantly higher performance than DeepLoc 2.0. Specifically, DualLoc-ProtT5 outperformed DeepLoc 2.0 across all key evaluation metrics, achieving an accuracy of 0.8113, a micro-averaged F1 score of 0.8941, and a macro-averaged F1 score of 0.8175 (Table 3). Performance gains were consistently observed across all variants utilising the same PLMs, confirming the broad applicability of this strategy to multilabel classification tasks. In addition, fine-grained analyses revealed that DualLoc-ProtT5 achieved substantial improvements in signal peptide classification precision. Among all variants, this model achieved the highest performance in predicting MT (MCC: 0.9584) and SP (MCC: 0.9052), as well as five other signal types. Moreover, ProtBERT achieved the highest performance in the TM category (MCC: 0.7835), whereas ESM-2 achieved the highest performance in the NES category (MCC: 0.5682). Overall, this optimised architecture enhanced the model’s ability to accurately recognise common sorting signals.

**Table 3.** Performance Evaluation on the Sorting Signal Peptide Dataset.

|  | DeepLoc 2.0 |  | DualLoc |  |  |  |
| --- | --- | --- | --- | --- | --- | --- |
|  | ESM1b | ProtT5 | ProtBERT | ESM1b | ESM-2 | ProtT5 |
| Pred. Num. Labels (1.20) | - | - | $1.1839 \pm 0.0195$ | $1.2047 \pm 0.0223$ | $1.2258 \pm 0.0207$ | $1.2305 \pm 0.0187$ |
| Accuracy | $0.78 \pm 0.02$ | $0.79 \pm 0.03$ | $0.7472 \pm 0.0228$ | $0.7878 \pm 0.0189$ | $0.7986 \pm 0.0148$ | **$0.8113 \pm 0.0132$** |
| Jaccard | - | - | $0.7023 \pm 0.0167$ | $0.6985 \pm 0.0241$ | $0.6839 \pm 0.0139$ | **$0.7097 \pm 0.0123$** |
| MicroF1 | $0.87 \pm 0.01$ | $0.87 \pm 0.02$ | $0.8384 \pm 0.0114$ | $0.8872 \pm 0.0098$ | $0.8927 \pm 0.0105$ | **$0.8941 \pm 0.0098$** |
| MacroF1 | $0.80 \pm 0.02$ | $0.80 \pm 0.03$ | $0.7629 \pm 0.0192$ | $0.8091 \pm 0.0174$ | $0.8101 \pm 0.0153$ | **$0.8175 \pm 0.0142$** |
| **MCC per signal** |  |  |  |  |  |  |
| – SP | $0.89 \pm 0.03$ | $0.90 \pm 0.03$ | $0.8637 \pm 0.0138$ | $0.8937 \pm 0.0291$ | $0.8928 \pm 0.0122$ | **$0.9052 \pm 0.0101$** |
| – TM | $0.71 \pm 0.07$ | $0.66 \pm 0.05$ | **$0.7835 \pm 0.0573$** | $0.7425 \pm 0.0538$ | $0.7524 \pm 0.0537$ | $0.7629 \pm 0.0485$ |
| – MT | $0.93 \pm 0.02$ | $0.93 \pm 0.03$ | $0.9021 \pm 0.0062$ | $0.9413 \pm 0.0212$ | $0.9536 \pm 0.0069$ | **$0.9584 \pm 0.0057$** |
| – CH | $0.85 \pm 0.07$ | $0.86 \pm 0.09$ | $0.8185 \pm 0.0531$ | $0.8654 \pm 0.0553$ | $0.8723 \pm 0.0552$ | **$0.8849 \pm 0.0498$** |
| – TH | $0.86 \pm 0.08$ | $0.80 \pm 0.08$ | $0.8289 \pm 0.0642$ | $0.8724 \pm 0.0659$ | $0.8771 \pm 0.0633$ | **$0.8895 \pm 0.0621$** |
| – NLS | $0.65 \pm 0.06$ | $0.66 \pm 0.01$ | $0.6027 \pm 0.0441$ | $0.6618 \pm 0.0473$ | $0.6814 \pm 0.0484$ | **$0.6976 \pm 0.0443$** |
| – NES | $0.49 \pm 0.20$ | $0.46 \pm 0.17$ | $0.4532 \pm 0.1754$ | $0.5072 \pm 0.1764$ | **$0.5682 \pm 0.1623$** | $0.5537 \pm 0.1689$ |
| – PTS | $0.85 \pm 0.06$ | $0.90 \pm 0.05$ | $0.7981 \pm 0.0511$ | $0.8789 \pm 0.0497$ | $0.8829 \pm 0.0534$ | **$0.8956 \pm 0.0507$** |
| – GPI | $0.85 \pm 0.06$ | $0.86 \pm 0.06$ | $0.8032 \pm 0.0523$ | $0.8815 \pm 0.0592$ | $0.8498 \pm 0.0519$ | **$0.8615 \pm 0.0482$** |

The best results are highlighted in bold.

### Associations Between Sorting Signals Classification and Attention Mechanism

DualLoc surpassed DeepLoc 2.0 in extracting and interpreting sorting signal features. Kullback–Leibler (KL) divergence was used to quantify the spatial consistency between attention distributions and true signal positions (Table 4). The results indicated that DualLoc achieved low KL divergence scores across five of nine signal types, with reductions ranging from 0.5% to 6.5%. These findings confirmed that the model’s attention mechanisms can precisely align with regions of biological signals. Fine-grained performance analyses further confirmed that DualLoc achieved higher classification accuracy across most sorting signals. Among all variants, DualLoc-ProtT5 exhibited the highest performance in SP (KL divergence: 0.9867), CH (KL divergence: 0.2887), and NES (KL divergence: 3.6774). In addition, DualLoc-ESM-1b achieved the lowest KL divergence for NLS (KL divergence: 2.5890) and GPI (KL divergence: 1.4904).

**Table 4.** KL Divergence of Attention Distributions in Sorting Signal Classification.

| Signal | DeepLoc 2.0 |  | DualLoc |  |  |  |
| --- | --- | --- | --- | --- | --- | --- |
|  | ESM1b | ProtT5 | ProtBERT | ESM1b | ESM-2 | ProtT5 |
| – SP | 1.04 ± 0.91 | 0.99 ± 0.86 | 1.0486 ± 0.8921 | 1.0254 ± 0.8973 | 1.0032 ± 0.8465 | **0.9867 ± 0.8799** |
| – TM | 1.13 ± 1.14 | 1.12 ± 1.03 | 1.1843 ± 1.0547 | 1.1122 ± 1.0188 | **1.0698 ± 0.9764** | 1.0901 ± 1.0299 |
| – MT | 0.51 ± 0.54 | 0.50 ± 0.48 | 0.5827 ± 0.5078 | 0.5086 ± 0.4821 | **0.5031 ± 0.4472** | 0.5729 ± 0.4915 |
| – CH | 0.32 ± 0.52 | 0.31 ± 0.31 | 0.3321 ± 0.3815 | 0.3035 ± 0.3152 | 0.2914 ± 0.2897 | **0.2887 ± 0.3068** |
| – TH | 0.19 ± 0.29 | 0.24 ± 0.16 | 0.2537 ± 0.1693 | 0.2087 ± 0.1729 | **0.1735 ± 0.1428** | 0.2430 ± 0.1590 |
| – NLS | 2.63 ± 1.52 | 2.60 ± 1.32 | 2.8975 ± 1.4029 | **2.5890 ± 1.3521** | 2.7538 ± 1.2796 | 2.8005 ± 1.3325 |
| – NES | 4.04 ± 1.51 | 3.88 ± 1.44 | 3.9012 ± 1.4765 | 3.7213 ± 1.4276 | 3.6929 ± 1.3987 | **3.6774 ± 1.4463** |
| – PTS | 0.85 ± 1.29 | **0.72 ± 1.05** | 0.8516 ± 1.0638 | 0.8671 ± 1.0374 | 0.8019 ± 1.0195 | 0.8585 ± 1.0504 |
| – GPI | 1.59 ± 0.73 | 1.85 ± 0.47 | 1.8548 ± 0.4825 | **1.4904 ± 0.4912** | 1.4936 ± 0.4512 | 1.8421 ± 0.4771 |

The best results are highlighted in bold. Lower values (↓) indicate superior performance.

Moreover, DualLoc-ESM-2 achieved the highest performance in the TM (KL divergence: 1.0698), MT (KL divergence: 0.5031), and TH (KL divergence: 0.1735) categories.

## Discussion

This study demonstrates that DualLoc, fine-tuning dual Transformer-based PLMs, significantly enhances protein subcellular localization prediction. Evaluations of Prot-BERT (bidirectional encoder), ESM-2 (RoBERTa variant for coevolutionary features), and ProtT5 (encoder-decoder with span masking) confirmed that full-parameter dual-model fine-tunings substantially outperform lightweight single-model approaches such as DeepLoc 2.0 [7], enabling deeper extraction of biologically relevant localization features [14, 24, 31].

This study further demonstrates that scaling laws [32, 33] for PLMs continue to hold for this downstream task. Our DualLoc framework integrates a pretrained PLM with a randomly initialized PLM in a cascaded architecture, augmented by attention and dropout layers. Importantly, this design increases model depth rather than width, enhancing representational capacity for complex sequence-to-function reasoning.

Swiss-Prot cross-validation demonstrated that DualLoc-ProtT5 outperformed the baselines, achieving improvements of 3.7% in accuracy (0.5872), 9.7% in micro-F1 (0.8271), and 12.1% in macro-F1 (0.7811). Key compartments—nucleus, extracellular space, and cell membrane—showed MCC gains of 0.13, 0.07, and 0.13, respectively.

Independent validation using HPA data confirmed the consistent superiority of full-parameter dual-model fine-tuning, underscoring its efficacy in capturing complex localization patterns.

Performance gains primarily derive from three factors. First, dual-model fine-tuning adapts pretrained representations to localization-specific features and extracts subtle contextual patterns that single-model approaches miss. Second, attention mechanisms dynamically prioritize key functional motifs. Third, dropout layers mitigate overfitting while enhancing cross-species generalizability. Sorting signals such as SP, MT, and PTS further increased interpretability, with reduced KL divergence confirming precise alignment of attention. Although applicable to ProtBERT, ESM-2, and ProtT5, computational-efficiency comparisons with DeepLoc 2.0 are limited by undisclosed training protocols.

Biological insights were well supported: Fig 3 confirmed Golgi-ER functional coupling (PMI = 0.25) in secretory pathways, aligning with literature [34]. Fig 4 showed strong signal-compartment specificity (80.2% →SP extracellular; 69.9%→TM membrane; about 100% NLS/MT→ nucleus/mitochondria). While the Sankey diagram maps the intricate routing pathways of multilocalized proteins, our performance metrics (Table 3) indicate that highly combinatorial scenarios—such as those driven by shuttling signals like NES or NLS—reduce decoding accuracy.

Future work should integrate LoRA to reduce the number of trainable parameters by *>*90% without compromising accuracy [35]. LoRA could halve GPU memory requirements [36], accelerate retraining on clinical datasets [37], and—coupled with data augmentation—improve convergence by 30% [38]. Structural data integration, like AlphaFold, and cross-species training would address current limitations in conformational signal modeling and taxonomic generalizability.

## Conclusion

This study demonstrates that full-parameter fine-tuning of dual PLMs—ProtBERT, ESM-2, and ProtT5—significantly outperforms lightweight single-PLM approaches for subcellular localization prediction. By updating all model parameters, our method extracts discriminative sequence features, enabling accurate classification across ten compartments. The DualLoc framework, integrating attention mechanisms and dropout layers, achieved comprehensive improvements over DeepLoc 2.0: accuracy (+5.4%), micro-F1 (+12.8%), macro-F1 (+9.2%), and key compartment MCC (+11.2–23.5%). Despite higher computational demands, full-parameter tuning yielded a 19.3% improvement in generalization on the independent HPA dataset, representing a favorable trade-off for multi-localization tasks. Future work will incorporate 3D structural information via graph neural networks and contrastive learning to enhance predictions for rare compartments, such as peroxisomes, and multi-localized proteins, thereby advancing applications in precision medicine and drug development.

## Code availability

The code and data supporting this study are openly accessible at https://github.com/NTOUBiomedicalAILAB/DualLoc.

## Supporting information

**S1 Table. Number of entries per category in the Swiss-Prot**.

**S2 Table. Number of entries per category in the independent HPA test set**.

**S3 Table. Counts for each sorting signal category**.

**S4 Table. DualLoc-ProtT5 Performance with five different dropouts**.

**S5 Table. AUROC scores on the Swiss-Prot Dataset**.

**S6 Table. AUPR scores on the Swiss-Prot Dataset**.

**S1 Fig. Impact of fine-tuning on ProtT5 feature representations**.

**(A)**PCA, UMAP, and t-SNE projections of protein sequence embeddings before fine-tuning. **(B)** Post-fine-tuning projections, demonstrating enhanced subcellular compartment clustering.

## Acknowledgments

We would like to thank Yan Cheng Lai for technical assistance and Drs. Ming Jing Hwang, Chia Yu Su, and Jia Ming Chang for their valuable feedback and insightful discussion.

## Author contributions

**Conceptualization:** Yan Guang Chen, Kuan Y. Chang.

**Data curation:** Yan Guang Chen.

**Formal analysis:** Yan Guang Chen, Wen-Yu Chung, Kuan Y. Chang.

**Funding acquisition:** Kuan Y. Chang.

**Investigation:** Yan Guang Chen, Kuan Y. Chang.

**Methodology:** Yan Guang Chen.

**Project administration:** Kuan Y. Chang.

**Resources:** Kuan Y. Chang.

**Software:** Yan Guang Chen.

**Supervision:** Wen-Yu Chung, Kuan Y. Chang.

**Validation:** Yan Guang Chen.

**Visualization:** Yan Guang Chen.

**Writing – original draft:** Yan Guang Chen, Kuan Y. Chang.

**Writing – review & editing:** Wen-Yu Chung, Kuan Y. Chang.

## References

1. Thul PJ, Åkesson L, Wiking M, Mahdessian D, Geladaki A, Ait Blal H, et al. A subcellular map of the human proteome. Science. 2017;356(6340):eaal3321.

2. Hung MC, Link W. Protein localization in disease and therapy. Journal of cell science. 2011;124(20):3381–92.

3. Lacoste J, Haghighi M, Haider S, Reno C, Lin ZY, Segal D, et al. Pervasive mislocalization of pathogenic coding variants underlying human disorders. Cell. 2024;187(23):6725–41.

4. Emanuelsson O, Brunak S, Von Heijne G, Nielsen H. Locating proteins in the cell using TargetP, SignalP and related tools. Nature protocols. 2007;2(4):953–71.

5. Imai K, Nakai K. Prediction of subcellular locations of proteins: where to proceed? Proteomics. 2010;10(22):3970–83.

6. Almagro Armenteros JJ, Sønderby CK, Sønderby SK, Nielsen H, Winther O. DeepLoc: prediction of protein subcellular localization using deep learning. Bioinformatics. 2017;33(21):3387–95.

7. Thumuluri V, Almagro Armenteros JJ, Johansen AR, Nielsen H, Winther O. DeepLoc 2.0: multi-label subcellular localization prediction using protein language models. Nucleic acids research. 2022;50(W1):W228–34.

8. Briesemeister S, Rahnenführer J, Kohlbacher O. YLoc—an interpretable web server for predicting subcellular localization. Nucleic acids research. 2010;38(suppl 2):W497–502.

9. Savojardo C, Martelli PL, Fariselli P, Profiti G, Casadio R. BUSCA: an integrative web server to predict subcellular localization of proteins. Nucleic acids research. 2018;46(W1):W459–66.

10. Gillani M, Pollastri G. Sclpred-ecl: subcellular localization prediction by deep n-to-1 convolutional neural networks. International Journal of Molecular Sciences. 2024;25(10):5440.

11. Zhang Y, Zheng L, You N, Hu W, Jiang W, Lu M, et al. LocPro: a deep learning-based prediction of protein subcellular localization for promoting multi-directional pharmaceutical research. Journal of Pharmaceutical Analysis. 2025;15(8):101255.

12. Vaswani A, Shazeer N, Parmar N, Uszkoreit J, Jones L, Gomez AN, et al. Attention is all you need. Advances in neural information processing systems. 2017;30.

13. Rives A, Meier J, Sercu T, Goyal S, Lin Z, Liu J, et al. Biological structure and function emerge from scaling unsupervised learning to 250 million protein sequences. Proceedings of the national academy of sciences. 2021;118(15):e2016239118.

14. Elnaggar A, Heinzinger M, Dallago C, Rehawi G, Wang Y, Jones L, et al. ProtTrans: toward understanding the language of life through self-supervised learning. IEEE transactions on pattern analysis and machine intelligence. 2021;44(10):7112–27.

15. Ødum MT, Teufel F, Thumuluri V, Almagro Armenteros JJ, Johansen AR, Winther O, et al. DeepLoc 2.1: multi-label membrane protein type prediction using protein language models. Nucleic Acids Research. 2024;52(W1):W215–20.

16. Shi H, Zhang X, Deng Q. ProStructNet: Integration of Protein Sequence and Structure for the Prediction of Multi-label Subcellular Localization. In: International Conference on Intelligent Computing. Springer; 2024. p. 326–36.

17. Jumper J, Evans R, Pritzel A, Green T, Figurnov M, Ronneberger O, et al. Highly accurate protein structure prediction with AlphaFold. Nature. 2021;596(7873):583–9.

18. Lin Z, Akin H, Rao R, Hie B, Zhu Z, Lu W, et al. Evolutionary-scale prediction of atomic-level protein structure with a language model. Science. 2023;379(6637):1123–30.

19. Luo Z, Wang R, Sun Y, Liu J, Chen Z, Zhang YJ. Interpretable feature extraction and dimensionality reduction in ESM2 for protein localization prediction. Briefings in Bioinformatics. 2024;25(2):bbad534.

20. Schmirler R, Heinzinger M, Rost B. Fine-tuning protein language models boosts predictions across diverse tasks. Nature Communications. 2024;15(1):7407.

21. Yuan GH, Li J, Yang Z, Chen YQ, Yuan Z, Chen T, et al. Deep generative model for protein subcellular localization prediction. Briefings in Bioinformatics. 2025;26(2):bbaf152.

22. Zeng GH, Zhu XZ, Yang HR, Liang YJ, Zhai YJ, Xu YY. Knowledge-enhanced protein subcellular localization prediction from 3D fluorescence microscope images. Bioinformatics. 2025;41(6):btaf331.

23. UniProt: the universal protein knowledgebase in 2021. Nucleic acids research. 2021;49(D1):D480–9.

24. Cui J, Yang S, Yi L, Xi Q, Yang D, Zuo Y. Recent advances in deep learning for protein-protein interaction: a review. BioData Mining. 2025;18(1):43.

25. Ling X, Li Z, Wang Y, You Z. Transformers in Protein: A Survey. arXiv preprint 250520098. 2025.

26. Refaeilzadeh P, Tang L, Liu H. Cross-validation. In: Liu L, Özsu MT, editors. Encyclopedia of Database Systems. Boston, MA: Springer US; 2009. p. 532–8. doi:10.1007/978-0-387-39940-9565.

27. Fano RM, Hawkins D. Transmission of information: A statistical theory of communications. American Journal of Physics. 1961;29(11):793–4.

28. Pearson K. LIII. On lines and planes of closest fit to systems of points in space. The London, Edinburgh, and Dublin philosophical magazine and journal of science. 1901;2(11):559–72.

29. McInnes L, Healy J, Melville J. Umap: Uniform manifold approximation and projection for dimension reduction. arXiv preprint 180203426. 2018.

30. Van der Maaten L, Hinton G. Visualizing data using t-SNE. Journal of machine learning research. 2008;9(11).

31. Gillani M, Pollastri G. Protein subcellular localization prediction tools. Computational and Structural Biotechnology Journal. 2024;23:1796–807.

32. Kaplan J, McCandlish S, Henighan T, Brown TB, Chess B, Child R, et al. Scaling laws for neural language models. arXiv preprint 200108361. 2020.

33. Rae JW, Borgeaud S, Cai T, Millican K, Hoffmann J, Song F, et al. Scaling language models: Methods, analysis & insights from training gopher. arXiv preprint 211211446. 2021.

34. Moremen KW, Tiemeyer M, Nairn AV. Vertebrate protein glycosylation: diversity, synthesis and function. Nature reviews Molecular cell biology. 2012;13(7):448–62.

35. Hu EJ, Shen Y, Wallis P, Allen-Zhu Z, Li Y, Wang S, et al. Lora: Low-rank adaptation of large language models. Iclr. 2022;1(2):3.

36. Liao X, Wang C, Zhou S, Hu J, Zheng H, Gao J. Dynamic adaptation of lora fine-tuning for efficient and task-specific optimization of large language models. In: Proceedings of the 2025 International Conference on Artificial Intelligence and Computational Intelligence; 2025. p. 120–5.

37. Saadat A, Fellay J. Fine-tuning protein language models to understand the functional impact of missense variants. Computational and Structural Biotechnology Journal. 2025;27:2199–207.

38. Sledzieski S, Kshirsagar M, Baek M, Dodhia R, Lavista Ferres J, Berger B. Democratizing protein language models with parameter-efficient fine-tuning. Proceedings of the National Academy of Sciences. 2024;121(26):e2405840121.

